# Mid-level feature differences underlie early animacy and object size distinctions: Evidence from EEG decoding

**DOI:** 10.1101/2022.01.12.475180

**Authors:** Ruosi Wang, Daniel Janini, Talia Konkle

## Abstract

Responses to visually-presented objects along the cortical surface of the human brain have a large-scale organization reflecting the broad categorical divisions of animacy and object size. Mounting evidence indicates that this topographical organization is driven by differences between objects in mid-level perceptual features. With regard to the timing of neural responses, images of objects quickly evoke neural responses with decodable information about animacy and object size, but are mid-level features sufficient to evoke these rapid neural responses? Or is slower iterative neural processing required to untangle information about animacy and object size from mid-level features? To answer this question, we used electroencephalography (EEG) to measure human neural responses to images of objects and their texform counterparts – unrecognizable images which preserve some mid-level feature information about texture and coarse form. We found that texform images evoked neural responses with early decodable information about both animacy and real-world size, as early as responses evoked by original images. Further, successful cross-decoding indicates that both texform and original images evoke information about animacy and size through a common underlying neural basis. Broadly, these results indicate that the visual system contains a mid-level feature bank carrying linearly decodable information on animacy and size, which can be rapidly activated without requiring explicit recognition or protracted temporal processing.

## 1 Introduction

The ventral visual stream contains extensive information about different object categories, with a large-scale spatial organization of response preferences characterized by the broad categories of animacy and object size (Grill-Spector and Weiner, 2014; Thorat et al., 2019; Konkle and Oliva, 2012; Konkle and Caramazza, 2013; Julian et al., 2017). Classic understanding of the ventral stream posits a hierarchical series of processing stages, en route to a more conceptual format that ultimately abstracts away from perceptual information (Mahon et al., 2009; Proklova et al., 2016, e.g. for review see Peelen and Downing, 2017). However, mounting evidence has revealed that the broad categorical distinctions of the ventral stream are driven by more primitive perceptual differences among “mid-level features” of texture, shape, and curvature (Long et al., 2016, 2017, 2018; Baldassi et al., 2013; Yue et al., 2020; Jozwik et al., 2016). On this emerging account of visual system processing, the ventral stream represents objects in a rich mid-level feature bank, from which more categorical distinctions can be extracted (e.g. with linear read-out).

Strong evidence for this mid-level feature bank account comes from recent work by Long et al. (2018) investigating brain responses to a new stimulus class called “texforms” (Long et al., 2016, 2017, 2018; **Figure 1A**). Texform images are created using a texture-synthesis algorithm (Freeman and Simoncelli, 2011) which preserves some mid-level feature information related to the texture and coarse form of the original depicted objects, while obscuring higher-level shape features like clear contours and explicit shape information. Empirically, people cannot identify what these are at the basic level (e.g. as a ‘cat’). Long et al. (2018) found that texform images evoked extensive responses along the entire ventral visual cortex with a similar large-scale organization as evoked by original, recognizable images. For example, zones of cortex responding more strongly to original animals also responded more to texformed animals. However, given that fMRI data obscure temporal information, there are a number of possible accounts of these large-scale activations. Thus, in the present study we examined the time-evolving signatures of visual system processing to ask when there is information about animacy and size in neural responses to texform images relative to their original counterparts.

**Figure 1:**
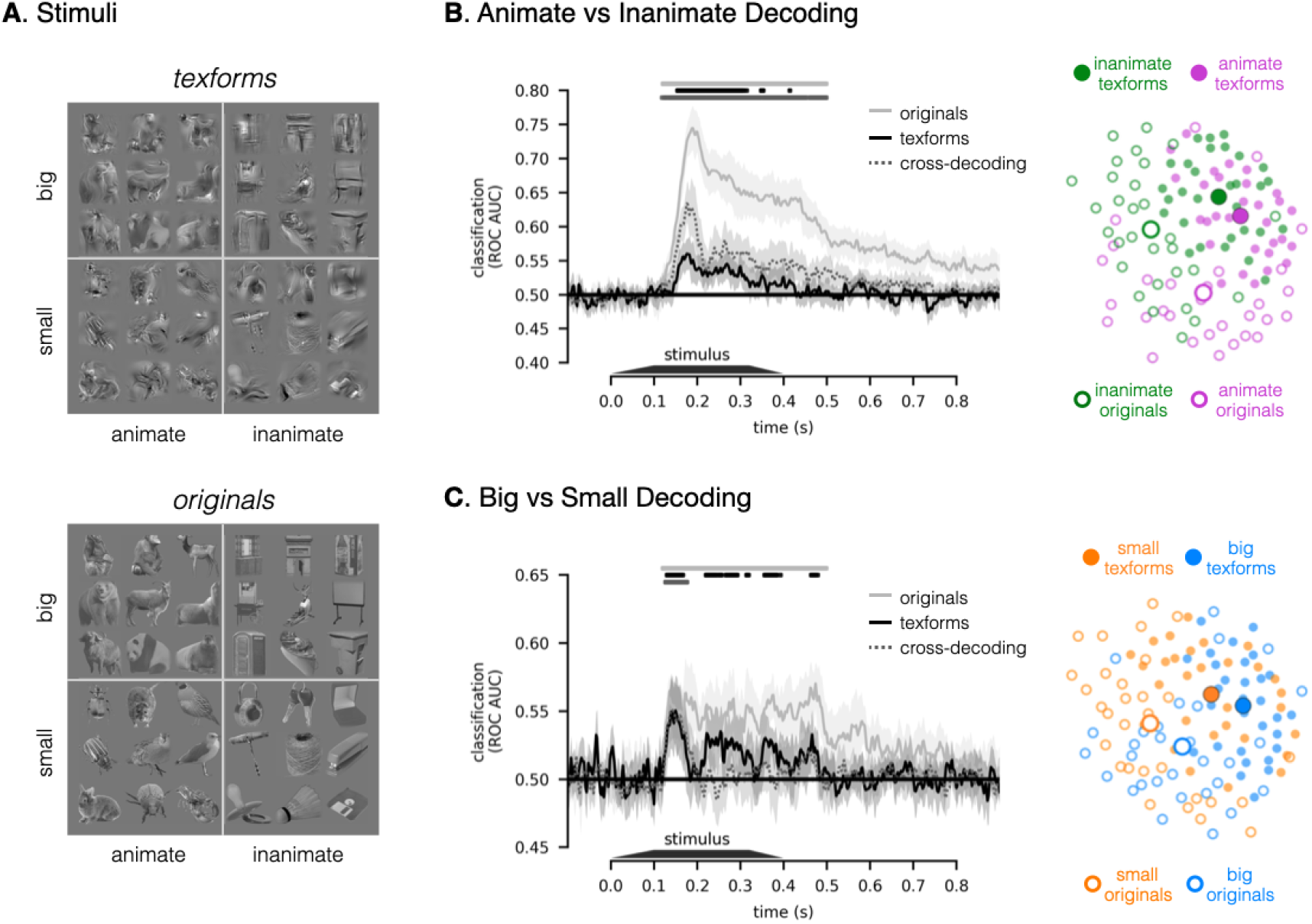
Stimuli and decoding results. ***A***, Example stimulus images. Each of the four conditions (animacy × size) included 15 exemplars, yielding 60 unrecognizable texforms (upper) and their 60 original counterparts (lower). See **Figure S1** for the full stimulus set. ***B***, Time course of animate vs. inanimate decoding. Classification accuracy is plotted along the y-axis, as a function of time (x-axis), for original (solid sliver lines), texform (solid black lines), and texform-to-original cross-decoding (dashed gray lines). Significant time points are depicted with horizontal lines above the time courses in the corresponding color (*p* < .05, one-sided signed-rank test, FDR corrected in the time window of interest, 100-500 ms). The shaded region indicates a 95% confidence interval. Adjacent to this axis is a multi-dimensional scaling (MDS) visualization, with a 2-D projection of the pairwise distances in the neural responses to each image from the peak animacy cross-decoding time (176 ms). ***C***, Time course for big vs. small decoding, as in ***B***. Adjacent MDS plot reflects a 2D projection of the neural similarity structure measured at the peak size cross-decoding time (140 ms).

According to the mid-level feature bank account, rapid feedforward activations of the ventral stream reflect sensitivity to mid-level featural distinctions which directly carry information about animacy and object size. A strong temporal prediction of this account is that animacy and object size information emerge early in the time-evolving responses, with comparable timing for texform and original formats. Indeed, EEG/MEG decoding studies measuring responses to intact pictures of objects have found that information about the animacy and real-world size can be decoded relatively early in the time course of processing (Carlson et al., 2013; Cichy et al., 2014; Kaneshiro et al., 2015; Ritchie et al., 2015; Grootswagers et al., 2017b,a; Khaligh-Razavi et al., 2018). Further, neurophysiological studies in non-human primates also have found that within 100 ms of stimulus onset, information about the animacy of the presented images can be decoded from the population structure of neural responses in V4 and IT (Cauchoix et al., 2016). Early decoding performance of these high-level properties in original images is consistent with a more primitive underlying format–though this inference is not required by the data.

An alternate temporal prediction is that neural responses to texforms will show more gradual emergence of animacy and object size information, increasing steadily over hundreds of milliseconds. This pattern of data might emerge if texforms contain only very subtle feature differences related to animacy and object size which are not linearly decodable in an initial feed-forward pass. These subtle differences may trigger later stages of processing which can reformat and amplify the visual input through more iterative processing steps, so that animacy and object size information is evident in the structure of the responses at later time points. Indeed, Grootswagers et al. (2019) recently argued for this possibility. They measured responses to texform and original images with EEG, using a rapid presentation design in which they varied the presentation speed of the stimuli. Considering neural responses to original images, they found that animacy and size information could be robustly decoded with presentation rates up to 30Hz. However, considering neural responses to texform images, they found that animacy could only be decoded at the slowest rate (5Hz), and size information was not decodable at all. Based on these results, they argued that texforms can elicit animacy signatures, but only given sufficient processing time, and that perhaps higher order visual areas are required to further “untangle” these features into linearly separable categorical organizations (DiCarlo and Cox, 2007).

Here we also measured EEG responses to both original and texform images depicting animate and inanimate objects of big and small real-world sizes. However, we used a standard event-related paradigm, allowing us to probe the structure of the neural responses without additional effects of forward and backward masking. To anticipate, we found that both animacy and size information could be decoded from EEG responses to texforms, as early in the time-evolving responses as evoked by original recognizable images. Moreover, we found that classifiers trained on neural responses to texform images were able to predict the animacy and size of responses to original images, indicating that these two image formats reflect animacy and object size information through a common representational basis. Broadly, our results thus support the view that mid-level feature differences contain signatures of animacy and object size which are available early in the visual processing stream.

## 2 Materials and methods

The experimental data and reproducible pre-processing/analysis scripts can be found at osf.io/mxrge.

### 2.1 Participants

Participants (n = 19) with normal or corrected-to-normal vision were recruited at the Harvard University community (mean age = 27.5, range: 20-42; 13 female; one left-handed). This sample size was decided by previous similar studies using EEG decoding (Grootswagers et al., 2017a; Bae and Luck, 2018). All participants provided informed consent and received course credits or financial compensation. We excluded one participant (female) from further analyses due to excessive movements and self-reports of discomfort during the experiment. All procedures were approved by the Institutional Review Board at Harvard University.

### 2.2 Stimuli and Tasks

The stimulus set consisted of 120 total images with 60 recognizable images of 15 big animals, 15 big objects, 15 small animals, and 15 small objects and their texform counterparts (**Figure 1A**, see **Figure S1** for the full stimulus set), which were created by undergoing a scrambling process (Freeman and Simoncelli, 2011). These stimulus images reflect a subset of the full stimulus set of Long et al. (2018), which consisted of 240 images. See Long et al. (2018) for detailed descriptions of stimulus generation.

Stimuli were presented on a 13-inch LCD monitor (1024 × 768 pixels; refresh rate = 60 Hz) at a viewing distance of around 60 cm with a visual angle of 12°, using MATLAB and Psychophysics Toolbox extensions (Brainard, 1997). A bullseye-like fixation remained present at the center of the screen at all times. At the start of each trial, an image was shown for a 400 ms stimulus presentation. In the first 100 ms of this time, the image was linearly faded in, and in the last 83.3 ms of this time, the image was linearly faded out. We made this choice based on the reasoning that it might reduce the abrupt onset and offset signals that might mask later-stage processing. At image offset, there was a 600 ms blank period before the subsequent trial began.

We instructed the participants to view the stimuli images attentively while undergoing EEG recording. Each run contained 240 trials (5.32 mins), with each stimulus exemplar appearing four times in a run. Participants first completed six runs of this protocol in which they saw texform stimuli, followed by six runs with original stimuli. To minimize artifacts, we included a 1.5-second “blinking period” every five trials. During this period, the fixation dot turned green to signal the participants that they were encouraged to blink. They were asked to refrain from blinking for the rest of the time.

### 2.3 EEG Recording and Preprocessing

Continuous EEG was recorded from 32 Ag/AgCI electrodes mounted on an elastic cap (EasyCap) and amplified by a Brain Products ActiCHamp system (Brain Vision) ^1^. The following scalps sites were used: FP1, FP2, F3, F4, FC1, FC2, Cz, C3, C4, CP1, CP2, CP5, CP6, P3, P4, P7, P8, POz, PO3, PO4, PO7, PO8, Oz, O1, O2, Iz, I3, I4. This montage was arranged according to the 10-20 system with some modifications. Specifically, three frontal electrodes were rearranged to have more electrodes over the posterior occipital pole (Long et al., 2017). Another two sites, T7 and T8, were also obtained but not used due to the noisy data. The horizontal electrooculogram (HEOG) was measured using electrodes positioned at the external ocular canthi to monitor horizontal eye movements. The vertical electrooculogram (VEOG) was measured at electrode FP1 to detect eye blinks. All scalp electrodes were online referenced to the average of both mastoids and digitized at a rate of 500 Hz.

We conducted EEG data preprocessing and analysis using the MNE-python package (Gramfort et al., 2014; https://mne-tools.github.io). First, portions of EEG containing excessive muscle movements were identified by visual inspection and removed. Continuous signals were then bandpass filtered with cutoff frequencies of 0.01 Hz and 100 Hz. In the next step, we applied ICA for each participant to identify and remove components associated with eye blinks or horizontal eye movements. The ICA-corrected data was segmented into 1000 ms epochs from – 100 ms to 900 ms relative to the stimulus onset and baselined to pre-stimulus periods. Finally, automated artifact rejection was employed to drop and repair bad epochs using the code package Autoreject (Jas et al., 2017) with default parameters.

Following these preprocessing steps, participants had on average 1373 trials (SD = 77) for texform stimuli and 1358 trials (SD = 85) for original stimuli, with no significant difference between these two stimulus types (*t*_(17)_ = .97, *p* = .35, paired *t*-test). The number of trials did not differ across conditions (big animals, big objects, small animals, and small objects) for either original (*F*_(3,51)_ = .96, *p* = .42, ANOVA) or texform (*F*_(3,51)_ = 1.12, *p* = .35, ANOVA) stimuli. We also conducted the main analyses without ICA and auto-reject procedures in place and obtained the same patterns of results.

### 2.4 Decoding Analysis

#### 2.4.1 Category-level decoding

A Linear Discriminant Analysis (LDA) classifier was trained to discriminate animate versus inanimate objects based on neural activation patterns across scalp electrodes, at each time point. The classifier was implemented with scikit-learn (Pedregosa et al., 2011) with default parameters.

We conducted decoding analysis on super-trials averaged across multiple trials rather than on single-trial data. This procedure is included because previous studies showed that averaging across several trials can improve the signal-to-noise ratio (Isik et al., 2014; Grootswagers et al., 2017b; Bae and Luck, 2018). In particular, six super-trials were computed for each stimulus exemplar by averaging over two to four trials since the numbers of trials varied across different stimuli after automatic artifact rejection. The number of averaged trials was determined by the recommendation of Grootswagers et al. (2017b). This procedure yielded 360 super-trials for recognizable stimuli (e.g., 180 animate / 180 inanimate) and 360 super-trials for texform stimuli. In addition, we also conducted data analysis without applying super-trial averaging and observed the same pattern of results.

To ensure the classifier can generalize to new stimuli, we used a 5-fold cross-validation procedure (Carlson et al., 2013). In each fold, the super-trials for 24 animate stimuli and 24 inanimate stimuli were used to train the classifier, which was then tested on the super-trials from the remaining 6 animate stimuli and 6 inanimate stimuli. For each fold, we measured the area under the curve of the receiver-operating characteristic (AUC ROC) which reflects an aggregate measure of performance across all possible classification thresholds.

To ensure the robustness of this AUC ROC estimate, we iterated the above procedure 20 times to minimize the idiosyncrasies in super-trial averaging and 5-fold stratified splits. After completing all iterations of cross-validation, the final decoding performance was computed as the average of the 100 decoding attempts (5 folds × 20 iterations).

#### 2.4.2 Cross-decoding

A similar decoding procedure was followed for the cross-decoding analyses, but trained on one stimulus type and tested it on the other. For example, in one fold, the classifier was trained using super-trials from 24 animate exemplars and 24 inanimate exemplars in their texform format. Critically, this classifier was then tested with super-trials from the remaining 6 animate and 6 inanimate exemplars in their recognizable form. We also conducted cross-decoding in the opposite direction (training on recognizable originals, testing on texforms) and reported in the **Figure S2**.

To create a graphical depiction of the similarity structure in the measured EEG responses, we used the following approach. First, electrode patterns were extracted for each object exemplar at each time point, yielding 60 conditions for recognizable images and 60 conditions for texform images. Next, we measured the multivariate noise-normalized (MNN) Euclidean distance (Guggenmos et al., 2018) between EEG patterns of all possible object pairs. Therefore, a 120 × 120 representational dissimilarity matrix was obtained for each participant at each time point. Finally, we used multidimensional scaling (MDS) to transform the group-averaged EEG RDM at the peak decoding time into a 2-dimensional space. Note that these plots are purely a supplementary visualization to provide a graphical intuition of the reported decoding results (e.g. the main decoding analyses were not conducted in this 2-dimensional MDS space).

#### 2.4.3 Pairwise decoding

To determine the decodability of each object against others, we estimated the pairwise decoding performance of all pairs of objects for both original and texform images. LDA classifiers were trained and evaluated with ROC AUC metric via five iterations of cross-validation. On each iteration, we trained a classifier to discriminate between two objects on 80% of trials and tested on the held-out 20% of trials. Please note that no super-trial averaging was applied here because of the limited number of trials for each single object stimulus (original: 22.6 ±1.4; texform: 22.9 ±1.3). The final pairwise decoding performance at each time point was the average of all pairwise decoding results across all cross-validation attempts (1770 pairs × 5 iterations). For the sake of saving computation time, we down-sampled the EEG data with a decimation factor of two.

#### 2.4.4 Onset and peak latencies

The time of decoding onset was defined as the first time point with above-chance decoding (*p* < .01, uncorrected) for three consecutive time points (Carlson et al., 2013). Note that in this procedure, multiple comparisons are not applied so that the estimation of onset latency does not depend on the decoding performance of later time points. The time of peak decoding was defined as the time point with maximum performance within the time window of interest. In the case where there were multiple local maximums within the window, the first of those maximums was selected.

### 2.5 Statistical Testing

To examine whether the decoding performance was significantly above chance, we conducted one-sided Wilcoxon signed-rank tests across the time points in the time window of interest (100-500 ms). This non-parametric test does not make any assumptions about the shape of the data distribution. When comparing the performance of different conditions of interest, we used two-sided Wilcoxon signed-rank tests. The obtained results were then corrected by FDR correction across all tested time points (*p* < .05).

We assessed the median and confidence interval of the onset and peak latencies using bootstrap sampling with 5000 iterations (bootstrapping the pool of participants with replacement). To test the differences of onset and peak latencies, we estimated the *p*-values based on bootstrapped distributions. Such results were corrected for the number of comparisons using FDR correction with the significance level of *p* < .05.

## 3 RESULTS

### 3.1 Animate vs. Inanimate Decoding

First, we examined whether recognizable images of animate and inanimate objects evoked distinguishable spatial EEG patterns over time, as has been previously shown (e.g., Carlson et al., 2013; Ritchie et al., 2015; Khaligh-Razavi et al., 2018; Grootswagers et al., 2017a). **Figure 1B** (solid silver line) shows a plot of decoding accuracy as a function of time for original images. Consistent with previous work, we observed a robust ability to classify animacy information: the spatial topography of the elicited EEG responses to animate and inanimate recognizable images were distinguishable from each other (*ps* < .05, one-sided signed-rank test, FDR corrected), with significant onset at 126 ms (95%CI: 116-142 ms) and peak classification accuracy at 188 ms (95%CI: 184-200 ms).

Next, we investigated (i) whether unrecognizable *texform* images of animate and inanimate objects evoke distinct spatial EEG patterns, and if so, (ii) at what time these distinctions emerge relative to the recognizable image counterparts. The same classification analysis as above was performed but considering only responses to texform images (**Figure 1B**, solid black line). Animate and inanimate texforms elicited different EEG patterns, with an onset of significant decoding at 152 ms (95%CI: 106-164 ms) and an early classification peak at 176 ms (95%CI: 146-190 ms). The onset and peak latencies of decoding for texform images did not significantly differ from those for recognizable images (onset: *p* = .34, peak: *p* = .14, bootstrapping test, FDR corrected). Critically, animacy decoding did not emerge over several hundreds of milliseconds, as would be predicted if extra processing time was needed to extract and/or amplify animacy information from texform images. However, animacy decoding did have a lower accuracy for texforms in comparison to original images (non-independent peak decoding accuracy: original 74.46% vs. texform 56.01%, *p* < .001, two-sided signed-rank test). Overall, these results indicate that the mid-level feature content preserved in texform images contains *early* perceptual signatures of animacy information.

Are the features that support the animacy distinction in texforms the same as those supporting animacy decoding in original recognizable images? If this is the case, both texforms and original images should evoke the same topographical differences that distinguish between animate and inanimate objects. To test this possibility, we conducted a cross-decoding analysis in which we trained a classifier to discriminate EEG responses to animate versus inanimate *texform* images, and then tested the classifier on responses to animate and inanimate *original* images. To ensure that the classifier was generalizing to new examples, we did not include any of the original-counterpart images to the texforms used to train the classifier. As shown in **Figure 1B** (dashed gray line), we found that texform-trained classifiers could successfully classify whether a new recognizable object was animate or inanimate (*ps* < .05, one-sided signed-rank test, FDR corrected). Such successful decoding was also evident early (onset: 140 ms, 95%CI: 114-152 ms; peak: 176 ms, 95%CI: 174-192 ms), with no significant difference in time to original images (onset: *p* = .44, peak: *p* = .34, bootstrapping test, FDR corrected) or texform images (onset: *p* = .14, peak: *p* = .46, bootstrapping testing, FDR corrected). Moreover, we also observed similar results when conducting the cross-decoding in the opposite direction (**Figure S2**, training on recognizable originals and testing on texforms). Thus, the classification boundary between animate and inanimate texforms also separates the animate and inanimate recognizable images, demonstrating the activation patterns are similar between these image formats.

Moreover, we unexpectedly found that texform-trained classifiers could predict the animacy more accurately for recognizable images than for other texform images (non-independent peak decoding: texform-original 63.4% vs. texform-texform 56.0%, *p* <.001, two-tailed signed-rank test). How is this superior classification accuracy possible? One possibility is that the original images evoke more discriminable neural responses than texforms, while still sharing a common large-scale topographic decision boundary. Consistent with this possibility, **Figure 1B** (right) provides a graphical intuition for this explanation. This multidimensional scaling (MDS) plot visualizes the neural pattern similarity structure among the original images (open dots) and texform images (filled dots) at the peak cross-decoding time (176ms), such that items with similar neural response patterns are nearby in the plot. Note that there is a general separation between animates (purple dots) and inanimates (green dots) across both texforms and originals. Further, the texform images (filled dots) are closer to each other; in contrast, recognizable images (open dots) are more distinctive and farther apart in this visualization.

Thus, this visualization helps provide an intuition for how original images can be classified more accurately than texform images by a texform-trained classifier.

A second piece of evidence also supports the interpretation that the original images evoke more separable, distinctive neural responses than those evoked by texforms. Specifically, we estimated the discriminability of responses at the item-level, estimating the average pairwise decoding accuracy over all pairs of items. **Figure 2** shows that pairwise decoding accuracy is significantly higher for original images than for texform images (*ps* < .05, two-sided signed-rank test, FDR corrected). Thus, we reason that, to the degree that both texforms and original images evoke similar patterns of neural responses that share a common decision boundary, the original images should be more easily classifiable because of their more distinctive evoked brain responses. In this way, this cross-decoding result provides strong evidence that the differences of elicited spatial topography that reflect animacy distinction in texforms are highly compatible with the distinguishing differences between recognizable animals and objects.

**Figure 2:**
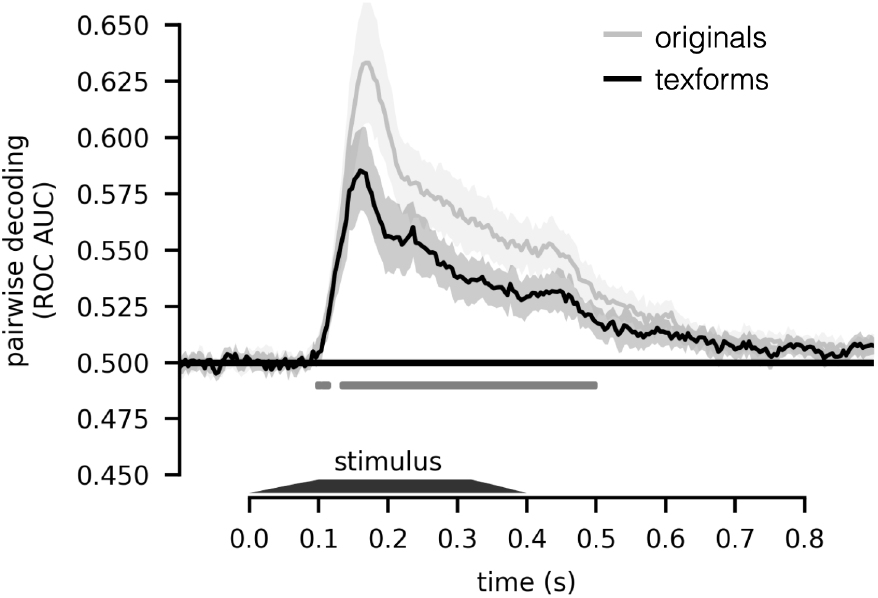
Time course of pairwise decoding. Pairwise decoding performance averaged across all object pairs is plotted along the y-axis, as a function of time (x-axis), for originals (silver line) and texforms (black line); shaded region indicates 95%CI. Time points with significant difference between original and texform stimuli are depicted below the time courses (two-sided signed-rank test, *p* < .05, FDR corrected in the time window of interest, 100-500 ms).

### 3.2 Big vs. Small Object Decoding

Next, we examined evoked differences between big and small entities, first by collapsing across the animate and inanimate dimension. Overall, the results reveal a similar pattern of results but with weaker overall decoding accuracy, plotted in **Figure 1C** (left). There was a significant difference between the elicited EEG response patterns to original images depicting big objects and small objects (*ps* < .05, one-sided signed-rank test, FDR corrected), as well as for texform images (*ps* < .05, one-sided signed-rank test, FDR corrected). Critical to the hypotheses under consideration, the timing of this emerging size distinction was also early in the response: neither decoding onsets nor decoding peaks for texform responses (onset: 130 ms, 95%CI: 120-246 ms; peak: 150 ms, 95%CI: 114-162 ms) were significantly different from those for original images (onset: 120 ms, 95%CI: 114-132 ms, *p* = .38; peak: 174 ms, 95%CI: 110-194 ms, *p* = .88, bootstrapping test, FDR corrected). Further, we found significant cross-decoding evident in classifiers trained on texform images and tested on original images (*ps* < .05, one-sided signed-rank test, FDR corrected), also evident early in time (onset: 124 ms, 95%CI: 120-128 ms; peak: 140 ms, 95%CI: 130-172 ms), with no significant difference in timing to original images (onset: *p* = .38, peak: *p* = .88, bootstrapping test, FDR corrected) or to texform images (onset: *p* = .38; peak: *p* = .88, bootstrapping test, FDR corrected). In summary, the above results demonstrate systematic (albeit weak) differences in neural responses to texformed versions of big and small images, evident early in the time course of processing, with compatible EEG response structure as evoked by original images.

We next considered size decoding separately for animals and object domains, motivated by previous work with fMRI by Konkle and Caramazza (2013). In particular, the spatial activations of ventral visual cortex exhibit three large-scale cortical zones preferentially responding to big inanimate objects, small inanimate objects, and animals (of both sizes). That is, there were similar spatial activation patterns for big and small animals (Konkle and Caramazza, 2013). Thus, we next examined the degree to which this “tripartite” signature was also apparent in the decoding of EEG responses. Given these previous findings from fMRI, we expected size decoding to be stronger among inanimate objects than among animals.

The results are shown in **Figure** 3. We found that size information was decodable from responses evoked by inanimate objects, and by animate objects, for both originals and texforms (all *ps* < .05, one-sided signed-rank test, FDR corrected). However, size decoding from responses to animal images was actually stronger than size decoding from responses to object images, contrary to what we expected (*ps* < .05, two-sided signed-rank test, FDR corrected). Note that this pattern of results held in both texforms and originals images. Considering the time course of this size decoding, responses to big vs small animals show an earlier and more rapid rise in their classifiability, whereas responses to big vs small inanimate objects show a slower and more gradual separability.

**Figure 3:**
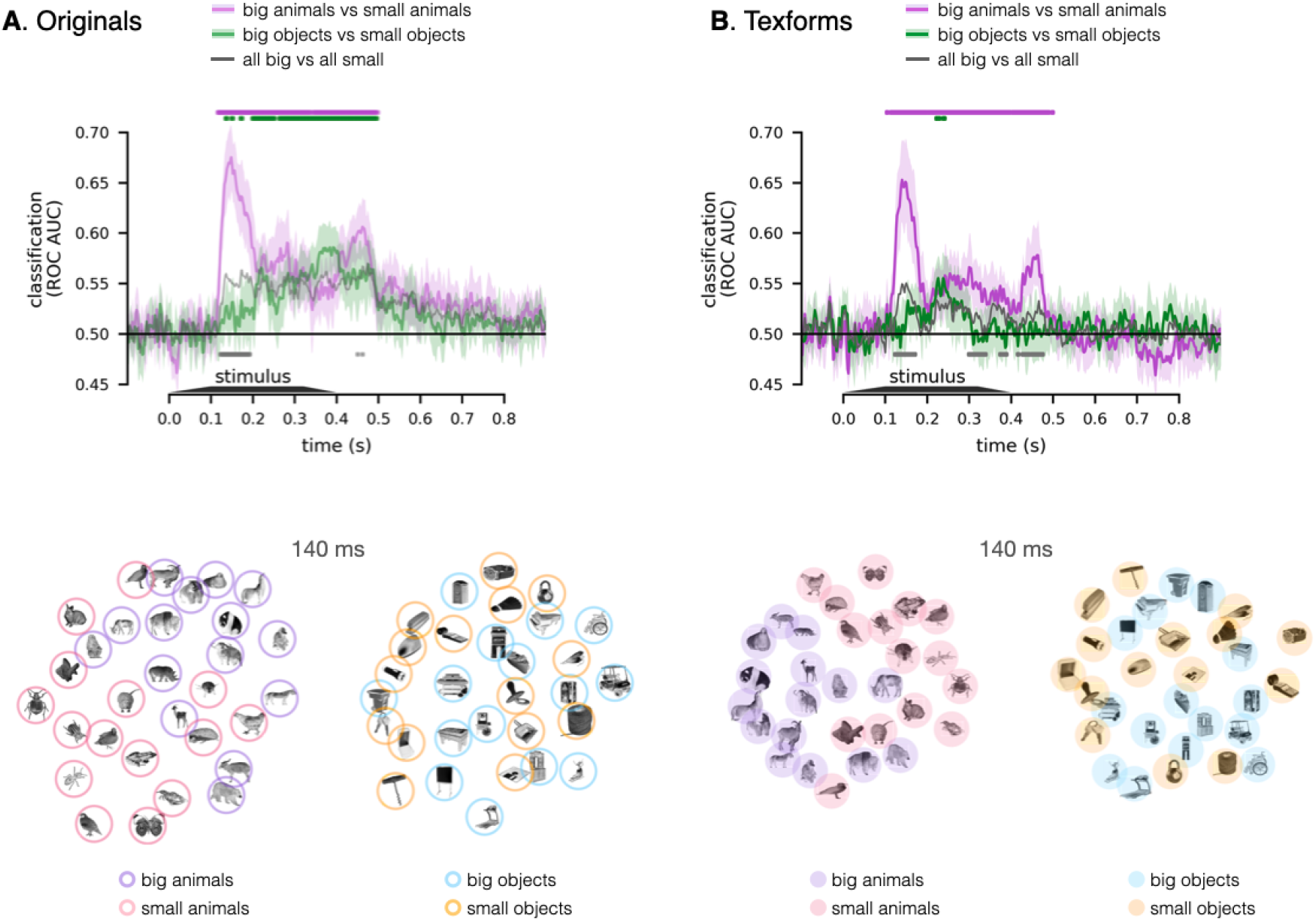
Decoding size among animate or inanimate objects only, for (**A**) original images and (**B**) texform images. In both plots (upper), classification accuracy (y-axis) is plotted as a function of time (x-axis). Purple line: big animals vs. small animals. Green line: big objects vs. small objects. Gray line: the combined classification for animates and inanimates is plotted for reference. Time points with significant decoding are depicted above the time courses (*ps* < .05, one-sided signed-rank test, FDR corrected in the time window of interest, 100-500 ms), and time points with significant difference between original and texform stimuli are depicted below the time courses (*ps* < .05, two-sided signed-rank test, FDR corrected). Blow the line plots are multi-dimensional scaling (MDS) visualizations, with a 2-D projection of the pairwise distances of the neural responses to animate objects only or inanimate objects only, examined at the peak cross-decoding time (140ms).

Using multidimensional scaling, we visualized the EEG pattern similarity structure among animals and among inanimate objects, separately for original and texform images (**Figure** 3). Specifically, we visualized the similarity structure evident at 140 ms, when the texform-to-original cross-decoding showed peak performance. In line with the decoding results, this visualization shows that the separation between big and small objects are clearer for animate objects in comparison to inanimate objects (for both original and texform images). Thus, these EEG size decoding results reveal a notable difference between the scalp-electrode response patterns over time and the large-scale cortical activation patterns along the ventral pathway that aggregated over time. We speculate on the underlying causes of these patterns of data in the General Discussion.

## 4 DISCUSSION

Here we employed multivariate EEG decoding to examine whether, and when, the visual system is sensitive to mid-level feature differences related to the broad distinctions of animacy and real-world size. We used a well established stimulus set that includes recognizable images of big and small animals and objects, as well as their unrecognizable “texform” counterparts (Long et al., 2016, 2017, 2018). We found that: (1) neural responses measured by EEG to texform images contained early information about animacy and size, as evident by above-chance decoding accuracy. (2) This broad categorical information was decodable from evoked responses to texforms at a similar time as from evoked responses to recognizable original images. (3) And, the time-evolving activation patterns were similar between these image formats, as evident by significant cross-decoding, suggesting a common underlying basis. Broadly, these EEG results indicate that the visual system contains an extensive mid-level feature bank, with early sensitivity to mid-level feature differences underlying animacy and size distinctions.

These patterns of data, and our subsequent interpretations, offer a different perspective than recent work by Grootswagers et al. (2019). Specifically, Grootswagers et al. also explored if animacy and size could be decoded from texform images, but they employed a fast image presentation paradigm in which the presentation rate was varied from 5Hz-60Hz–differing from our slow event-related design. In their data, texforms elicited brain response structure with weaker decoding of animacy information than recognizable objects, and only at the slowest presentation rate. Based on these results, they proposed that additional processing time in higher order visual areas is required to further “untangle” the mid-level feature differences evident in texforms into linearly separable categorical organizations (c.f. DiCarlo and Cox, 2007). In contrast, we propose that no further “untangling” is required for animacy and object size information to emerge.

To reconcile our findings with Grootswagers et al. (2019), we offer the following account. We propose that the visual system contains a mid-level feature bank which carries linearly decodable information on animacy and size. Texforms and original images rapidly activate this feature bank in a primarily feedforward processing sweep, enabling early decoding accuracy. Crucially, texforms don’t elicit the same magnitude of feature activation as original images–this is evident in our data by their generally lower decoding accuracy, both at the category and item-level, and is also found in neuroimaging results (Long et al., 2018). Thus, even though responses to texforms rapidly activate the same features as original images, these responses may be more likely to be extinguished under conditions of masking. This proposal accounts for the similarities between texforms and originals seen in our study, and posits increased susceptibility to masking during rapid texform presentation.

How do the current real-world size decoding results relate to previous fMRI work? Specifically, Konkle and Caramazza (2013) found that big and small object images evoked a large-scale organization of responses across the cortical surface, while big vs small animals had similar response topographies. We expected that EEG decoding accuracy would also reflect this tripartite organization, but that is not what we found. We can rule out the possibility that the distinction between big and small animals was driven by the detection of recognizable eyes or frontal faces, because this result was also evident in the texform images which lack clear facial features. One possibility, invited by the time-course of decoding, is that the neural populations which distinguish between big and small animals are only engaged early and transiently, and their responses may not be evident in slower aggregated responses of fMRI. This spatial-temporal hypothesis may be possible to explore through fMRI-MEG fusion (Cichy et al., 2016; Khaligh-Razavi et al., 2018), ECOG, or neural recordings in monkey populations. More generally, these results highlight the need for a deeper exploration of the convergences and discrepancies between the spatial similarity structure of neural activation patterns over EEG electrodes, and BOLD-estimated activations over cortical voxels.

One limitation of the present study is that the number of stimuli employed was relatively limited (N=60, 15 per animacy-size combination), leaving open the possibility that these randomly selected exemplars may not be fully representative of the broader categories they were sampled from. In our analyses, we leveraged cross-validation methods which require predicting animacy and size in held-out stimuli, mitigating this concern with an analytical approach. Further, this work joins a growing set of results showing the tight links between original and texformed counterparts in perceptual processes (e.g. Long et al., 2016, 2017; Chen et al., 2021), and more generally between mid-level feature distinctions and broader categorical distinction (Groen et al., 2017). Overall, this work provides clear support for the claim that early visual processes operating over mid-level features contain information about the broad categorical distinctions of animacy and object size.

## Author contributions

R.W., D.J. and T.K designed research. R.W. and D.J. performed research. R.W.analyzed data. R.W., D.J., and T.K. interpreted the results. R.W. and T.K, wrote the first draft of the paper. R.W., D.J., and T.K. edited the paper. R.W. and D.J. contributed unpublished analytic tools.

## Acknowledgements

We thank Aylin Kallmayer and Hrag Pailian for their help during the experiments and data collection. This work was supported by NSF CAREER BCS-1942438 (T.K.).

## Footnotes

1 Early in piloting, we tested this paradigm both with our newer 64-channel EEG system and with a custom channel configuration with more electrodes over the visual cortex. These equipment changes did not yield any differences in the overall pattern of our pilot data. Thus, we went to the 32-channel system because the set-up time was much shorter, which enabled us to increase the power per subject within the limited duration of an EEG experimental session.

