## Supplemental Figures for "Mid-level feature differences underlie early animacy and object size distinctions: Evidence from EEG decoding"

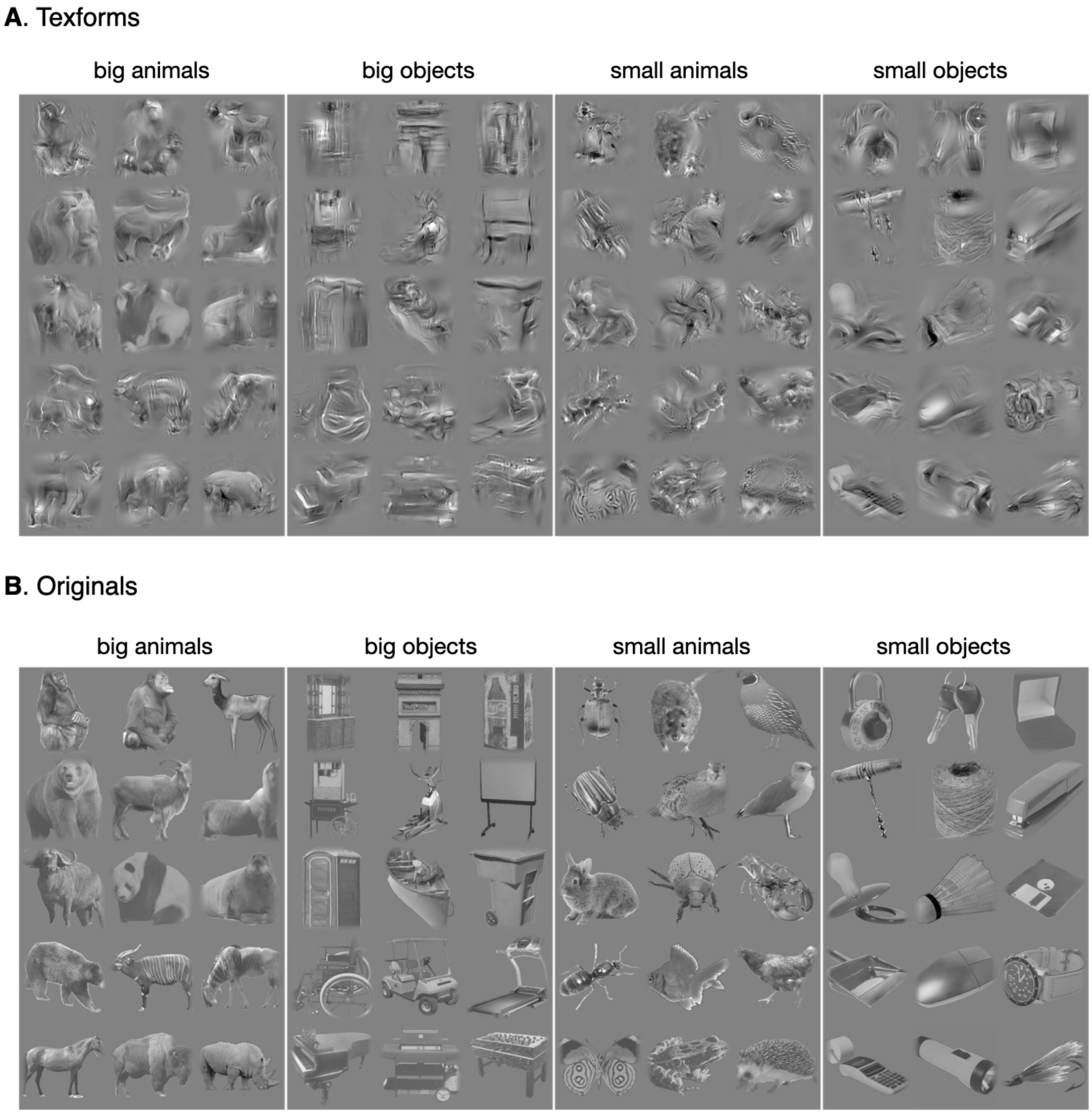


**Figure S 1:** Stimulus sets for (**A**) original and (**B**) texform images. Fifteen total exemplars were included for the 4 animacy x size conditions, yielding 60 unrecognizable texforms and their 60 original counterparts.

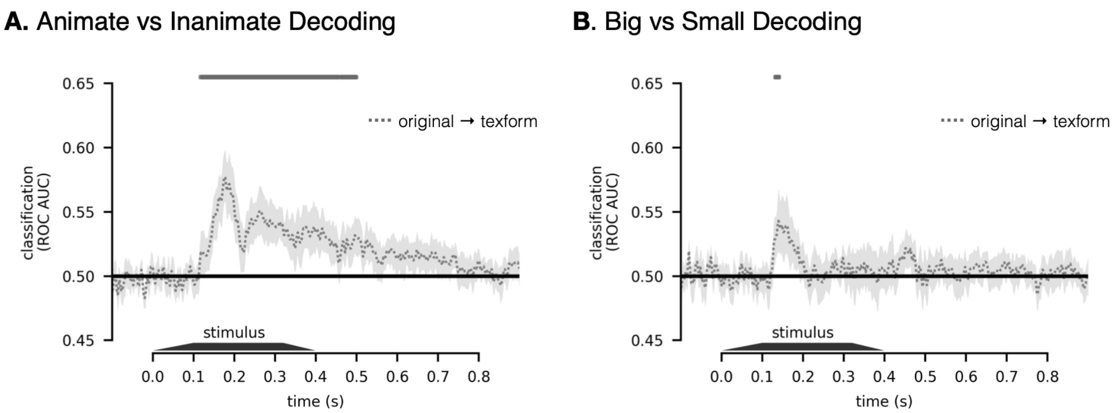


**Figure S 2:** Original-to-texform cross decoding results, for (**A**) animate vs. inanimate decoding and (**B**) big vs. small decoding. In both plots, classification accuracy (y-axis) is plotted as a function of time (x-axis). Significance, indicated above the plots, reflects one-sided signed rank test, $p < .05$, FDR corrected in the time window of interest, 100-500 ms).
